# Direct measurement of sub-kilobase chromatin structure reveals that linker histone H1 broadly compacts chromatin, with differential impact amongst epigenetic states

**DOI:** 10.64898/2025.12.19.695525

**Authors:** Hera Canaj, Irene Duba, Andres Mansisidor, Andrew Scortea, Ryan Johnson, Hugo Pinto, Arnold Ou, Nicole Pagane, Juhee Pae, Dmitry Fyodorov, Arthur I. Skoultchi, Viviana I. Risca

**Affiliations:** Laboratory of Genome Architecture and Dynamics, The Rockefeller University, New York, NY; Department of Cell Biology, Albert Einstein College of Medicine, New York, NY; Laboratory of Lymphocyte Dynamics, The Rockefeller University, New York, NY

## Abstract

Chromatin compaction by linker histone H1 family proteins is a long-standing model for transcriptional repression. However, the biophysical and conformational details of such compaction *in situ*, at the kilobase- and sub-kilobase length scale relevant to the activity of transcriptional regulatory elements, remain under debate. Rather than inferring such compaction from indirect measurements of features like DNA accessibility, we sought to directly probe sub-kilobase contacts between nearby nucleosomes. We developed an improved version of radiation-induced correlated cleavage with sequencing (RICC-seq), which we term RICC-seq 2.0, and used it in parallel with Micro-C to cross-validate our measurements of chromatin structure in both diverse cell types with different levels of linker histone and different levels of chromatin compaction, as well as a CRISPRi system for pan-H1 depletion. Using this system, we find that chromatin fiber de-compaction upon H1 depletion is global across the genome, reducing the contrast in inter-nucleosome contacts between acetylated chromatin and the rest of the genome. Surprisingly, this does not dramatically change higher-order chromatin organization such as nuclear compartments. Nevertheless, we observe a broad increase in accessibility at tens of thousands of sites and an increase in expression of over a thousand genes, which are enriched in polycomb repressive complex targets. Investigating the local chromatin compaction at upregulated genes as opposed to genes that do not change transcription, we observe that upregulated genes are not specifically de-compacted. Rather, our data support a model in which linker histone globally induces local compaction of nucleosome contacts and an increase in linker lengths, and repression by PRC1/2 is particularly dependent on these local features of chromatin architecture.

## Main Text

Compaction of the chromatin fiber has been invoked as a mechanism for transcriptional repression since the earliest investigations into the organization of chromatin fibers *in vivo* and *in vitro*^1–4^. *In situ*, microscopy shows that the density of silent heterochromatin is significantly higher than the density of transcriptionally active euchromatin^5,6^. *In vitro*, chromatin fibers with regularly spaced nucleosomes collapse into compact 30 nm-diameter structures, supporting the “30-nm fiber” model of transcriptional repression^7^. H1 linker histones are a family of proteins essential for development in metazoans, which bind the dyad positions of core nucleosomes via their globular domain and interact with the linker DNA entering and exiting the nucleosome via their unstructured, positively charged N-terminal and C-terminal domains ^8^. In a variety of chromatin reconstitution experiments, linker histones have been shown to promote the compaction of chromatin fibers or chromatin domains into higher density structures ^7,9–12^.

Although the simple 30-nm fiber model dominated the field for decades, efforts to assess *in vivo* or *in situ* chromatin fiber structure enabled by advances in electron microscopy and X-ray scattering did not identify the expected long-range regular higher-order helices posed by the 30-nm fiber model ^13–16^. Chromatin was therefore proposed to be an unstructured, liquid-like “polymer melt” of nucleosomes^3^. This updated model of chromatin as a liquid is consistent with the recently observed propensity of unmodified chromatin to form phase-separated liquid condensates^12,17^. However, even in condensates, local structural motifs of chromatin fibers dictated by nucleosome modifications and the geometry of inter-nucleosome stacking interactions dictated by linker DNA lengths and architectural proteins such as linker histones can modulate phase separation behavior^17^. For example, linker histones were shown to increase the density of chromatin condensates^12^, and sequencing-based methods for mapping local chromatin interactions, such as Micro-C and RICC-seq, found short-range zig-zag tetranucleosome folding signatures^18–20^. Super-resolution imaging found that chromatin fibers consist of small clusters of nucleosomes, termed “clutches”, the size of which is modulated by factors including linker histones^21^. In vitro FRET measurements and simulations both point to such clusters or tetranucleosome motifs being highly dynamic^22,23^.

A full understanding of chromatin fiber structure and behavior, and the regulation of its interactions with the proteins that carry out DNA-based processes, including DNA replication, transcription, and DNA repair, therefore requires us to reconcile the long-range disorder and the potential local order of chromatin. This is particularly important as the local interactions of nucleosomes determine the accessibility and binding affinity of individual loci to these proteins. For example, the spacing of nucleosomes, which, as a result of DNA’s helical nature and its relative stiffness on the length scale of typical inter-nucleosome linker lengths (∼30-70 bp), strongly determines the geometry of nearby nucleosome stacking, is tightly controlled by nucleosome remodelers, forming regular arrays in some areas of the genome^17,24–28^. Linker histones are one of the strongest determinants of nucleosome spacing, with high linker histone expression correlating with longer average linker lengths and a large nucleosome repeat length (NRL)^29^. Long linker segments can create boundaries between nucleosome interaction domains^30^. Structural studies show that chromatin modifiers (“writers”) can bridge nucleosomes as they propagate the histone modification from one nucleosome to another, either alone, like the polycomb repressive complex 2 (PRC2) or through dimerization^31,32^, as is the case for the H3K9me3-binding protein HP1. The efficiency of deposition of H3K27me3 by PRC2 *in vitro* is enhanced by compacted chromatin fibers^6,33^. On the other hand, the efficiency of the modification H3K36me2 by the methyltransferase NSD2 is inhibited by chromatin fiber compaction^6^.

We therefore sought to understand how local chromatin fiber compaction on the scale of a few nucleosomes—the length scale relevant to protein binding at regulatory regions on DNA—is regulated by linker histone in different chromatin contexts and what its consequences are for transcriptional repression. Locus-specific chromatin compaction *in situ* is very difficult to measure, and proxies for compaction such as DNA accessibility measured by ATAC-seq have historically been used instead. To address this challenge, we sought to use two complementary approaches that rely on different fundamental operating principles: Micro-C^18^ and radiation-induced correlated cleavage with sequencing (RICC-seq)^20^.

Micro-C, which relies on cell crosslinking, micrococcal nuclease (MNase) digestion of chromatin, and proximity ligation of DNA ends, can probe nucleosome-nucleosome contacts and chromatin organization on the kilobase scale, and differences in local contacts, such as a zig-zag-like signature in nucleosome contact probabilities, between yeast and mouse embryonic stem cells (mESCs) ^18^. This is *a priori* attractive as a potential measure of compaction, but uncertainties about artifacts caused by sequence and particularly by the accessibility bias of MNase cleavage make it difficult to determine whether the differential signal observed is due to compaction or accessibility.

This uncertainty motivated our use of an orthogonal method to Micro-C in order to validate results and gain more sensitivity to local changes in nucleosome contacts. RICC-seq^20^ relies on spatial clusters of DNA damage events, within a few nanometers of each other, that produce characteristic single-stranded DNA fragment lengths in irradiated cells. The peaks in the fragment length distribution (FLD) reflect the lengths of frequently occurring DNA loops spanning self-contact points that are simultaneously cleaved within a diameter of ∼8 nm. The primary peaks observed in RICC-seq FLDs from human fibroblasts correspond to a single DNA wrap around a nucleosome (∼78 nt), a full nucleosome unit (∼180 nt), and contacts between the DNA gyres of stacked alternating nucleosomes (∼270 nt and ∼360 nt). Using chromatin fiber simulations, we explored how the locations and strengths of these peaks vary with chromatin fiber geometry, indicating that RICC-seq FLDs have the potential to be sensitive to nucleosome spacing, nucleosomal DNA wrapping (which alters DNA entry/exit angles), and the strength of attractive interactions between nucleosomes. This indicated to us that RICC-seq should be able to detect the effects of linker histone H1 on local chromatin compaction, beyond what is already known about its effects on nucleosome spacing and linker lengths.

Before applying RICC-seq to this problem, we had to overcome its limitations: the protocol was long, requiring more than a week to complete, did not compensate for sample-to-sample variations in DNA fragment length capture bias to enable quantitative comparisons between different samples, and exhibited significant sequence bias in the final libraries. Here, we develop an optimized RICC-seq 2.0 protocol that solves these challenges, and apply it, together with Micro-C, to measure DNA-DNA contacts on the sub-kilobase length scale in cells with varying levels of H1 linker histones. We find that linker histone has a dramatic genome-wide effect on kilobase-scale chromatin compaction, and that changes in accessibility and transcription are concentrated in regions silenced by the polycomb repressive complexes (PRC1/2).

## Results

### RICC-seq 2.0 improves robustness, reduces sequence bias, and allows cross-sample comparison

In order to use RICC-seq to assess the chromatin fiber compaction effects of linker histone across cell types and perturbation conditions, we improved on the original RICC-seq method in several ways.

First, we addressed a challenge we encountered when scaling up the RICC-seq protocol to larger numbers of samples and conditions: library preparations would sometimes fail, yielding no peaks in the fragment length distribution. The original protocol used a high heat denaturation step to dissociate radiation-cleaved single-stranded DNA (ssDNA) fragments from the higher molecular weight genomic DNA prior to elution. We found that this heat-elution method caused the appearance of a large number of additional ssDNA breaks at heat-labile sites throughout the genome, overwhelming the DNA cleavage signal from the original radiation-induced breaks. These heat-labile sites have been previously documented as a product of DNA irradiation^34^. Small variations in the precise timing of heat denaturation would cause more or fewer of these breaks, leading to a lack of robustness in the RICC-seq protocol. Replacement of heat denaturation with high-pH (NaOH incubation) denaturation and avoidance of high heat (above 65°C) in subsequent library processing was sufficient to generate a more robust ssDNA elution and library preparation (Figure 1a-b).

**Figure 1.**
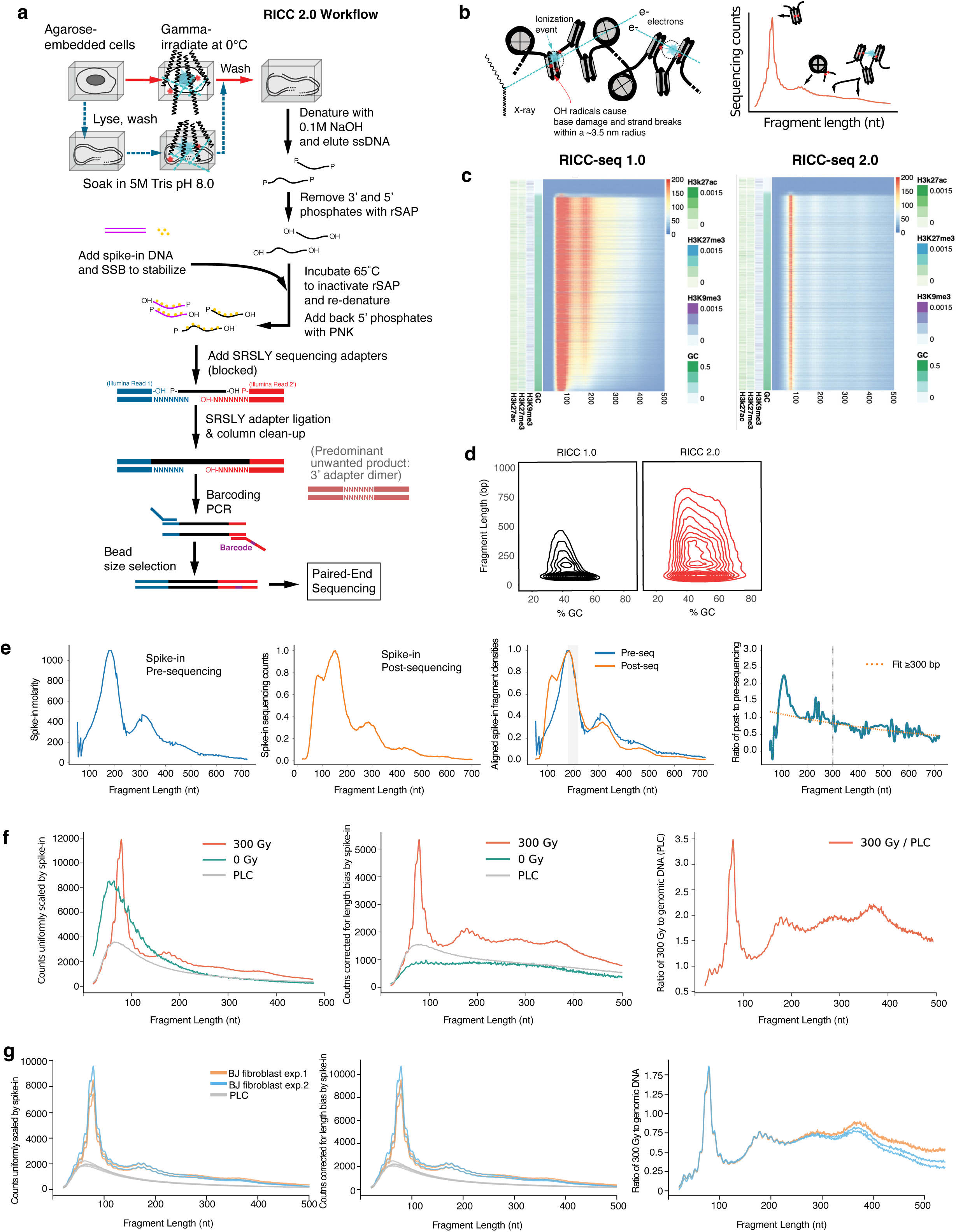
Improved RICC-seq 2.0 protocol reduces sequence bias, improves recovery of long high-GC fragments and allows quantitative comparison between samples. **a)** Schematic of the new RICC-seq protocol, incorporating the SRSLY ssDNA library preparation, including spike-in. **b)** Schematic depicting fragments produced by RICC-seq and the nucleosome-nucleosome contacts that generate the corresponding peak distributions. **c)** Fragment-length distribution plot over increasing genome %GC with fragments mapping to Roadmap Epigenomics ^40^ H3K27me3, H3K9me3, and H3K27ac peaks shown for RICC-seq 1.0 and 2.0 methods. **d)** Contour plot of fragment lengths captured and %GC of representative BJ sample from RICC-seq 1.0 and RICC-seq 2.0. **e)** MNase-digested fission yeast chromatin ladder is shown before sequencing as a Capillary electrophoresis (TapeStation) trace and after sequencing as a Fragment Length Distribution (FLD), both smoothed (5 nt rolling average) and aligned by the falloff of the mononucleosome peak. Ratio of post- to pre-sequencing distributions shown with exponential curve fit after 300bp which is extrapolated back and used to correct samples within an experiment. **f)** Representative BJ-5ta fibroblast sample with the respective corrections applied **g)** Multiple BJ fibroblast samples shown with the respective corrections applied n=2 biological replicates shown each with n=2 technical replicates.

Second, we addressed another challenge of using RICC-seq—the length and complexity of the protocol—which was partly due to the necessity for end-repair in agarose plugs and multiple gel-based size selections and amplifications to strike a balance between maintaining as much of the insert size distribution as possible while removing dimers of ligated sequencing adapters. To streamline sequencing adapter ligation to the eluted ssDNA, we used the Single Reaction Single-stranded LibrarY (SRSLY) protocol developed for ancient DNA and cell-free DNA sequencing, which uses single-strand binding protein (SSB) to stabilize and blocked adapters with random-heptamer splint overhangs to capture the end-repaired ssDNA fragments^35^. This allowed us to proceed from ligation to PCR without the need for size selection (Figure 1a).

Together, these changes produced a more robust RICC-seq 2.0 protocol that can capture ssDNA fragments from irradiated cells across a broader range of GC contents and fragment lengths (Figure 1c). In particular, RICC-seq 2.0 demonstrates an improved efficiency of capture for fragments that are both long and GC-rich (Figure 1d).

Third, because RICC-seq can be sensitive to the length bias introduced by sample handling and PCR, we developed a spike-in and normalization strategy to account for such sample-specific biases and allow us to quantitatively compare samples across experiments, cell types and perturbations. To create a “standard candle” library, we digested *Schizosaccharomyces pombe* chromatin with MNase into a nucleosome ladder, quantified its FLD using capillary electrophoresis prior to library preparation. Known quantities of this *S. pombe* spike-in were then added to RICC-seq libraries prior to SRSLY adapter ligation and library preparation (Figure 1a). Spike-in reads were computationally isolated to calculate their own FLD and a length bias correction factor was calculated by fitting an exponential to the ratio of the post-sequencing FLD and pre-sequencing FLD (Figure 1e). The absolute amount of spike-in was used to scale FLDs for comparisons between no-irradiation controls, irradiated genomic DNA, and irradiated cell samples (Figure 1f). The length-dependent correction factor (Figure 1e) was then used to correct length bias in RICC-seq FLDs (Figure 1f). Lastly, to enhance the contrast of peaks in the FLD over the background of DNA fragments caused by random, uncorrelated breaks, we calculated the ratio of the scaled and corrected FLD from irradiated cells to the scaled and corrected FLD from irradiated genomic DNA from the same cell sample (maintained in 0.5M Tris pH 8.0 as a quencher for radiation-induced radicals, approximating intracellular quenching) (Figure 1f). These procedures allowed us to directly compare replicates and different experimental samples that may have been subject to different length biases (Figure 1g).

### RICC-seq 2.0 is sensitive to chromatin compaction differences across species

The combination of a more robust protocol and spike-in normalization allowed us to apply RICC-seq 2.0 (Figure 1) to a broad range of cell samples, including budding yeast (Figure 2a). Budding yeast does not express a canonical member of the H1 linker histone family, but expresses Hho1p, which is homologous to the H5 linker histone found in chicken erythrocytes and binds nucleosomes ^36^. However, its expression level is much lower than mammalian linker histones: a ratio of 0.3 molecules per nucleosome ^29,36^. Budding yeast therefore has short inter-nucleosome linkers and a short NRL, and a largely open chromatin conformation with little clear distinction between euchromatin and heterochromatin as is found in metazoans. At the other extreme, the linker histone to nucleosome ratio rises even higher to ∼1.3 in the transcriptionally inactive nuclei of chicken erythrocytes, in which the primary histone variant is H5^29^. Chicken erythrocytes have been used as a model system for highly compacted chromatin fibers, as they represent one of the few cell types in which electron microscopy reveals structures resembling 30-nm diameter fibers, albeit more disordered ones than reconstituted in vitro ^5,9,37^.

**Figure 2.**
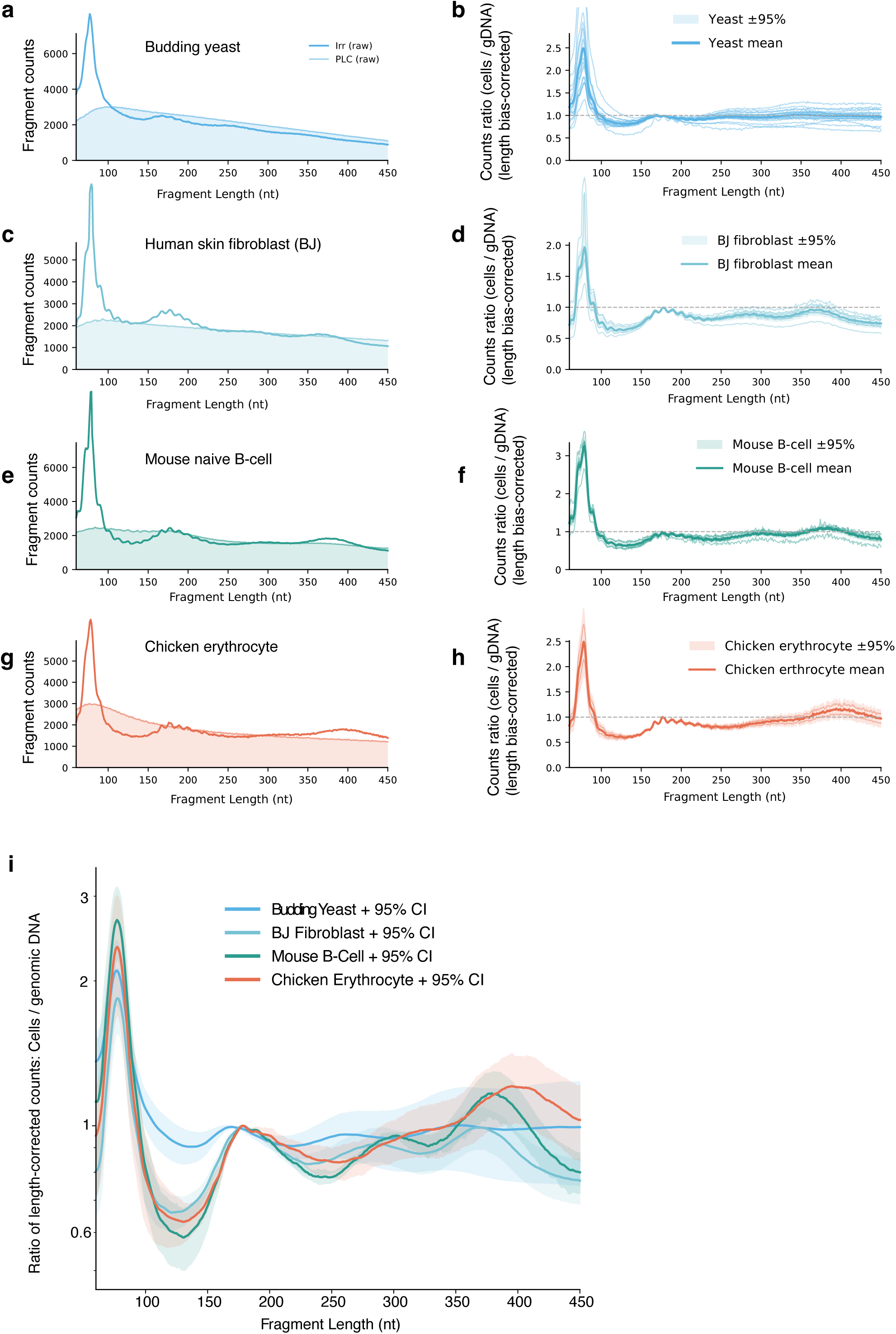
RICC-seq 2.0 applied across organisms with increasing H1 levels and chromatin compaction reveals an increase in nucleosome stacking contacts. **a,c,g,e)** Representative spike-in–scaled FLDs for each species before length-bias correction **b,d,h,f**) Replicate FLDs after length-bias correction with 95% CIs, max-normalized to the mononucleosome signal; condition means shown with 95% CI **i)** Per-organism corrected replicate FLDs averages shown. Chicken n=2 technical replicates, Mouse B cell n=2 technical replicates, BJ n=3 technical replicates of 2 biological replicates, yeast n=3 technical replicates of two biological replicates.

Seeking to understand the dynamic range of RICC-seq 2.0 FLDs as a function of varying compaction and linker histone levels, we applied it to four sample types: *S. cerevisiae,* human BJ-5ta fibroblasts, naïve mouse B cells, and chicken erythrocytes (Figure 2a-h). After correcting for length bias and calculating the ratio of the irradiated cell sample FLDs to their corresponding irradiated genomic DNA FLDs, we compared them directly (Figure 2i). We found two main effects. First, moving from budding yeast, with an average NRL of 163 bp, to BJ-5ta human fibroblasts, with an average NRL of 186 ^38^, mouse B cells with a NRL estimated to be ∼192 (based on human lymphoblastoid cells^28^), to chicken erythrocytes, with an average NRL of 212 bp ^29^, we observed a shift in the location of the higher-order contact (third and fourth) peaks of the RICC-seq FLD toward longer fragment lengths, consistent with the increase in FLD. Importantly, we also observed that the inter-nucleosome contact peaks were more prominent compared to the sub-nucleosomal (first) and mono-nucleosome (second) peak in cell types with more compact chromatin, such as human fibroblasts and to a greater extent, mouse B-cells. Chicken erythrocytes had the most extreme example, with a high fourth peak at ∼400 nt.

The correlation between the linker histone level and the inter-nucleosome stacking signal, which we interpret as a measure of local chromatin fiber compaction differences between cell types, motivated us to perturb the linker histone level in a well-characterized system to more precisely analyze its effects, context dependence and functional consequences.

### Linker histone depletion by CRISPRi leads to genome-wide reduction of nucleosome repeat length and loss of zig-zag alternating nucleosome contacts

Using a doxycycline-inducible dCas9 K562 cell line, we designed CRISPRi guides against the four H1 subtypes that are the most abundantly expressed in K562 cells: H1.2, H1.3, H1.4 an H1.5, as well as scrambled controls (Figure 3a). The guide RNAs were stably transfected. dCas9 induction for five days led to a reduction in the H1:nucleosome ratio from ∼0.75 to ∼0.2, as quantified by HPLC (Figure 3b-c). Over the five-day timeline of the experiment, cell doubling time was not qualitatively different, indicating maintenance of viability (Figure 3d).

**Figure 3.**
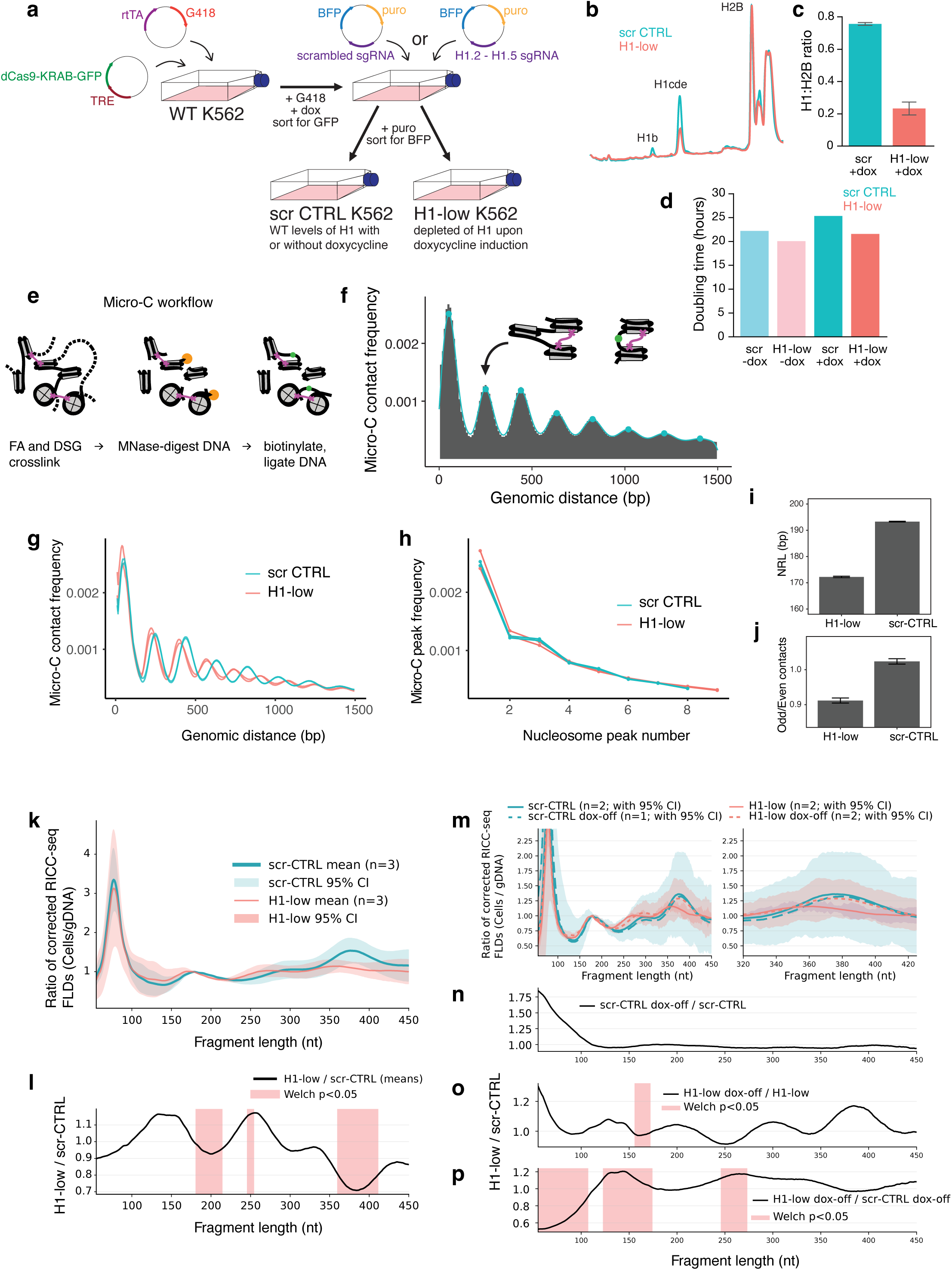
H1 depletion shortens nucleosome repeat length and reduces contacts associated with nucleosome-nucleosome stacking interactions, which reemerge upon wash-out. **a)** Experimental design for generating H1-low and scrambled control (scr-CTRL) cells, with five days doxycycline (dox) induction and matched 5 days dox-washout (rescue) conditions for induction of dCas9 in cells with constitutively expressed, stably transfected CRISPRi guides RNAs targeting H1.2, H1.3, H1.4, and H1.5 (H1-low) or scrambled guides (scr-CTRL). **b)** HPLC of H1 subtypes in H1-low cells relative to scr-CTRL. **c)** Quantification of HPLC of H1:H2B ratio depletion on day 5 of dox induction. **d)** Doubling time shown for samples before and after induction on dox for scr-CTRL (blue) and H1-low (pink). n=1 biological replicate. **e-f)** Schematic of Micro-C workflow and resulting short-range contact probability histogram. **g)** Micro-C contact frequency curve and **h)** maximum contacts for scr-CTRL and H1-low conditions **i)** nucleosome repeat length (NRL) quantification n=2, **j)** odd/even contact frequency peak maximum quantification of n=2 biological replicates shown. **k)** RICC-seq fragment length distribution in scr-CTRL and H1-low cells, spike-in–scaled, depth-matched within biological replicate, normalized to the 180 nt peak maximum, interpolated to a common 55–450 bp grid, and smoothed (10 nt rolling average). Shaded bands denote 95% CI (t-interval) across the relevant replicates n= 3 biological replicates shown. **l)** Ratio taken over scr-CTRL and H1-low; pink boxes mark contiguous Welch-significant runs (p<0.05; calculated within 5 nt windows) for the indicated pairwise comparison. **m)** Condition means across dox washout rescue experiment (scr-CTRL vs H1-low). n=2 H1 low with dox, n=2 H1low dox washout, n= 2 Scr with dox n=1 scr-CTRL dox washout. n: technical replicates. **n)** Ratio of technical replicate means from (m) ±95% CI for scr-CTRL, scr-CTRL dox-off. **o)** Ratio of technical replicate means from (m) ±95% CI for H1-low and H1-low dox-off. **p)** Ratio of technical replicate means from (m) ±95% CI for H1-low dox-off/scr-CTRL dox-off; shaded segments denote Welch test-significant runs (p<0.05; 5 nt windows).

We applied Micro-C (Figure 3e-f) to cells with CRISPRi-depleted H1 (H1-low) and control cells expressing scrambled CRISPR guides (scr-CTRL) at the 5-day time point. Due to potential differences in global accessibility upon reduction of linker histone and the sensitivity of Micro-C, as with other MNase-based assays, to the precise MNase concentration, we titrated and optimized the MNase concentration for each condition independently until similar chromatin digestion profiles were obtained, as assayed by capillary electrophoresis.

Analysis of the short-range (< 1.5 kb) Micro-C contact probability curves (Figure 3f) revealed that both scr-CTRL cells and H1-low cells exhibit a series of peaks corresponding to contacts between integer nucleosome steps proceeding down the fiber (N+1, N+2, …), with two main differences between the curves (Figure 3g-h). Most prominent is a shift in peak location corresponding to a drop in the NRL upon H1 depletion (Figure 3g,i). However, a second, more subtle but significant effect is a difference in the relative heights of the contact frequency peaks. While scr-CTRL cells exhibit a staircase-like pattern in which pairs of contact peaks are of similar height (N+2 and N+3, N+4 and N+5, …), this pattern was subdued and the peak heights approached a smoothly decreasing function in the H1-low cells (Figure 3h). We quantified this through the ratio of odd and even nucleosome contact probability peaks and found the effect to be significant across our biological replicates (Figure 3j).

Although matching the global MNase digestion profiles between samples should mitigate some of the accessibility biases of Micro-C on a genome-wide scale, concerns that the Micro-C results may not fully reflect local folding of the chromatin fiber nevertheless remain. To validate that our observed change in not only NRL but also nucleosome contact (and hence chromatin fiber folding) patterns are not an artifact of differential MNase digestion, we applied the RICC-seq 2.0 protocol to the same cells, using a *S. pombe* spike-in to normalize fragment length histograms between samples. RICC-seq does not rely on enzymatic digestion and should therefore not be influenced in the same way by changes in the accessibility to proteins. Its cleavage events are mediated by ionizing radiation that penetrates the whole nucleus and by highly diffusible species—primarily, hydroxyl radicals^20^. In genome-wide analysis, we found that the RICC-seq results corroborated our findings from Micro-C (Figure 3k-p). The inter-nucleosome peaks shift to lower fragment lengths in the H1-low RICC-seq FLD, indicating a lower NRL (Figure 3k), and the strength of the fourth peak, which was most strongly correlated with chromatin compaction and linker histone levels in our cross-species comparison (Figure 2), dropped significantly (Figure 3k-l). Smaller significant changes were also present in the second (mono-nucleosome) RICC-seq FLD peak (Figure 3k-l), but we do not draw a strong conclusion from this segment of the FLD because it exhibited more variability between biological replicates.

We then sought to determine to what extent these effects on the short-range chromatin fiber compaction evident in the RICC-seq data were a direct result of linker histone depletion, as opposed to indirect effects, such as from cell stress responses. We performed a washout experiment in which the H1-low and scr-CTRL cells were depleted of H1 for five days, as before, and then cultured in doxycycline-free media for five more days to allow for H1 levels to return. We found that the strength and location of the fourth peak returned upon dCas9 washout (Figure 3m-p). Overall, this led us to conclude that linker histone H1 has a direct effect on short-range stacking between alternating nucleosomes—nucleosome N to N+2 zig-zag contacts—in the context of intact chromatin.

### Short-range zig-zag stacking contrast between euchromatin and heterochromatin depends on linker histone levels

Next, we asked how the dependence of nucleosomal zig-zag contacts depend on the local epigenetic context. Segmenting the Micro-C contacts by overlap with histone mark ChIP-seq peaks—H3K27 acetylation to mark active promoters and enhancers, and H3K27 trimethylation and H3K9 trimethylation to mark the two primary types of heterochromatin—we found that there was a subtle change in the zig-zag signal of the first eight Micro-C contact peaks (Figure 4a-b). Quantitating the zig-zag signature using the odd-even peak height ratio, we found that there were differences in compaction between the three chromatin states, with H3K9me3 chromatin having the strongest zig-zag and H3K27me3 the weakest (Figure 4c). H1 depletion reduced the zig-zag signature such that the H3K27me3 heterochromatin in H1-low cells had a similar level to H3K27 acetylated chromatin in scr-CTRL cells (Figure 4c).

**Figure 4:**
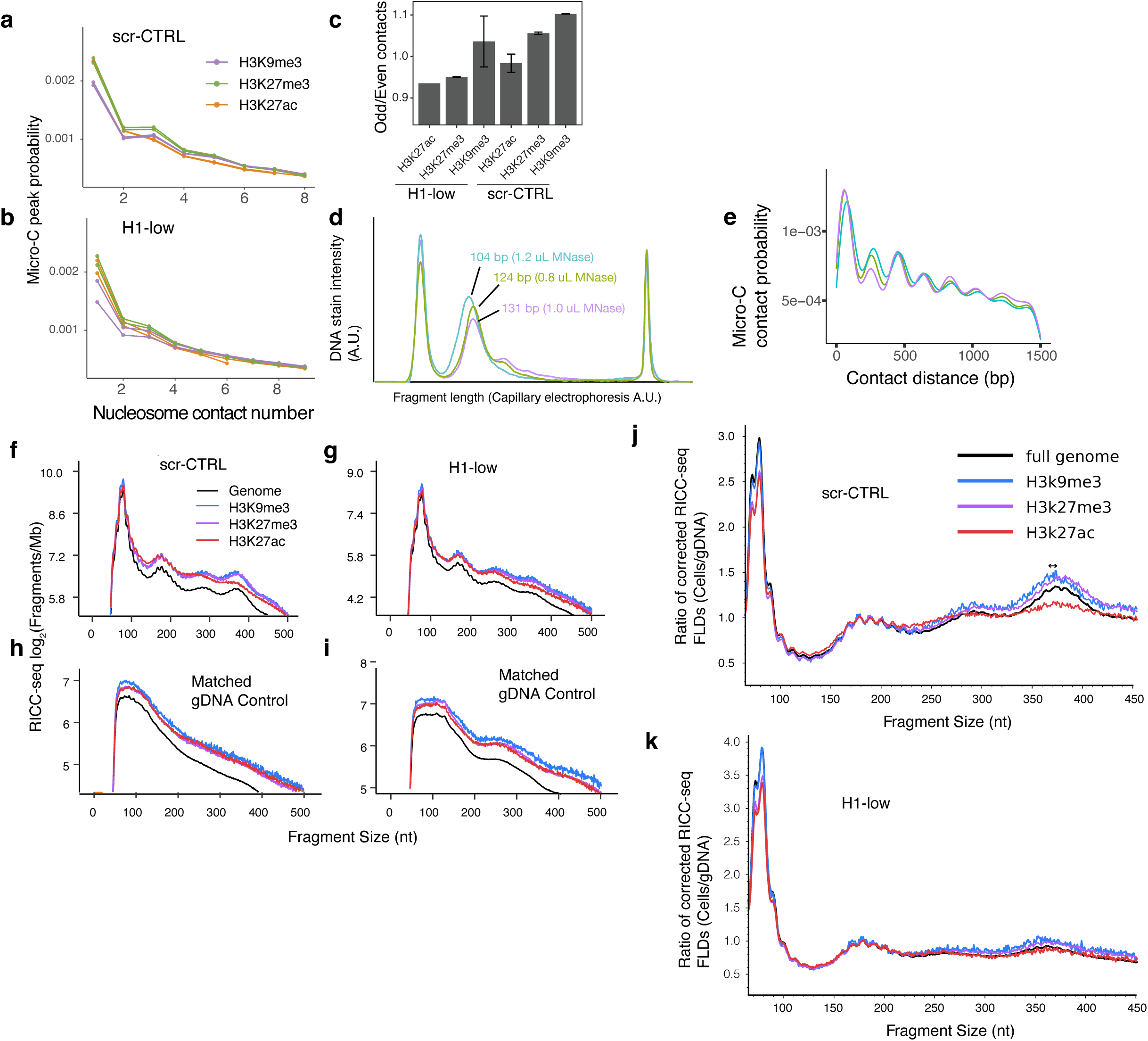
H1 depletion phenocopies compaction structure of active chromatin. Nucleosome-nucleosome contacts as measured by Micro-C in epigenetic regions defined by published WT K562 ChIP datasets in **a)** scr-CTRL **b)** and H1-low conditions ^40^. Two biological replicates shown each. **c)** Ratio of N/N+odd contacts and N/N+even nucleosome contacts in epigenetic state regions. Error bar: standard deviation between biological replicates, n=2. **d)** Capillary electrophoresis (TapeStation) traces of the fragment size distribution produced by MNase titration with different amounts of enzyme. The estimated fragment length of the mononucleosome peak is indicated. **e)** Micro-C contact probability curves for the libraries obtained from the MNase titration in **(d)**. **f-g)** RICC-seq FLDs from irradiated cells and matched gDNA controls in h-i) **j-k)** scr-CTRL cells and H1-low cells, subset by histone mark ^40^ gapped peaks. 1 biological replicate shown.

We cautiously interpret the zig-zag signature in short-range Micro-C contact data on a global level as a measure of short-range chromatin compaction because differences in the propensity for cleavage by MNase can be controlled at a global level by tuning the MNase concentration. However, artifacts caused by differential digestion by MNase cannot be mitigated if they occur between different sets of genomic loci within the same sample, as would be expected for heterochromatic *versus* euchromatic loci. Indeed, we observed that in libraries with different extents of digestion, as measured by the effective fragment size of the mononucleosome peak (Figure 4d), the zig-zag signature depended on the amount of digestion, with an inverse correlation between the strength of the zig-zag signature and the size of the mononucleosome fragment (Figure 4d-e). We therefore concluded that Micro-C is not a reliable measure of differences true chromatin compaction within the same sample, and validation of results by an orthogonal method is needed.

We analyzed our RICC-seq 2.0 data segmented by epigenetic state in the same mode, in order to determine whether the patterns of zig-zag contacts suggested by the Micro-C data could be orthogonally validated by a non-enzymatic method (Figure 4f-k). We monitored the irradiated genomic DNA (gDNA) control from both scr-CTRL and H1-low cells to ensure that the peak changes we observed were not driven by pre-existing DNA damage that could be differential between genomic loci (Figure 4h,i). Although we observed some weak peaks in the H1-low gDNA control consistent with small amounts of contaminating DNA fragments with damage between nucleosomes, they were not correlated with the irradiated cell peaks in a way that would explain the observed differences. To normalize against differences in the gDNA, we calculated the ratio between the length bias-corrected, epigenetic state-specific RICC-seq cell FLDs to the similarly corrected gDNA FLDs (Figure 4j-k). The results we observed partially agree with Micro-C data. Interpreting the strength of the fourth peak (∼330-420 nt) as a measure of the population-averaged stacking of alternating nucleosomes by zig-zag chromatin fiber compaction, we found that acetylated chromatin indeed does have very low compaction. However, the level of compaction between H3K27me3 heterochromatin and H3K9me3 heterochromatin appears similar by RICC-seq, as opposed to the higher H3K9me3 compaction suggested by Micro-C. The location of the fourth peak is shifted (Figure 4j, arrow) between H3K9me3 and H3K27me3 chromatin, consistent with the difference in NRL observed by Micro-C.

The difference in local chromatin fiber compaction between heterochromatic regions and acetylated euchromatic regions is consistent with the removal of linker histone H1 from acetylated chromatin causing unfolding of the fiber ^10,39^. We therefore compared the epigenetic state compaction landscape between scr-CTRL cells and the H1-low cells. We found that most of the difference in chromatin fiber compaction between epigenetic states in the scr-CTRL landscape is gone in H1-low cells (Figure 4g,k).

### Changes to long-range chromatin compartments, domains, and loops are minimal after five-day H1 depletion

The dramatic loss of local chromatin fiber compaction upon H1 depletion observed with short-range Micro-C curves and RICC-seq 2.0 motivated us to ask how this short-range decompaction relates to long-range chromosome folding features. We performed Hi-C to obtain a sensitive measure of long-range compartment changes. The large-scale (1 Mb resolution) balanced contact matrices appeared similar between scr-CTRL and H1-low cells (Figure 5a) and indeed, HiC-Rep analysis at 500 kb resolution showed that the difference between conditions was comparable to the difference between replicates (Figure 5b).

**Figure 5:**
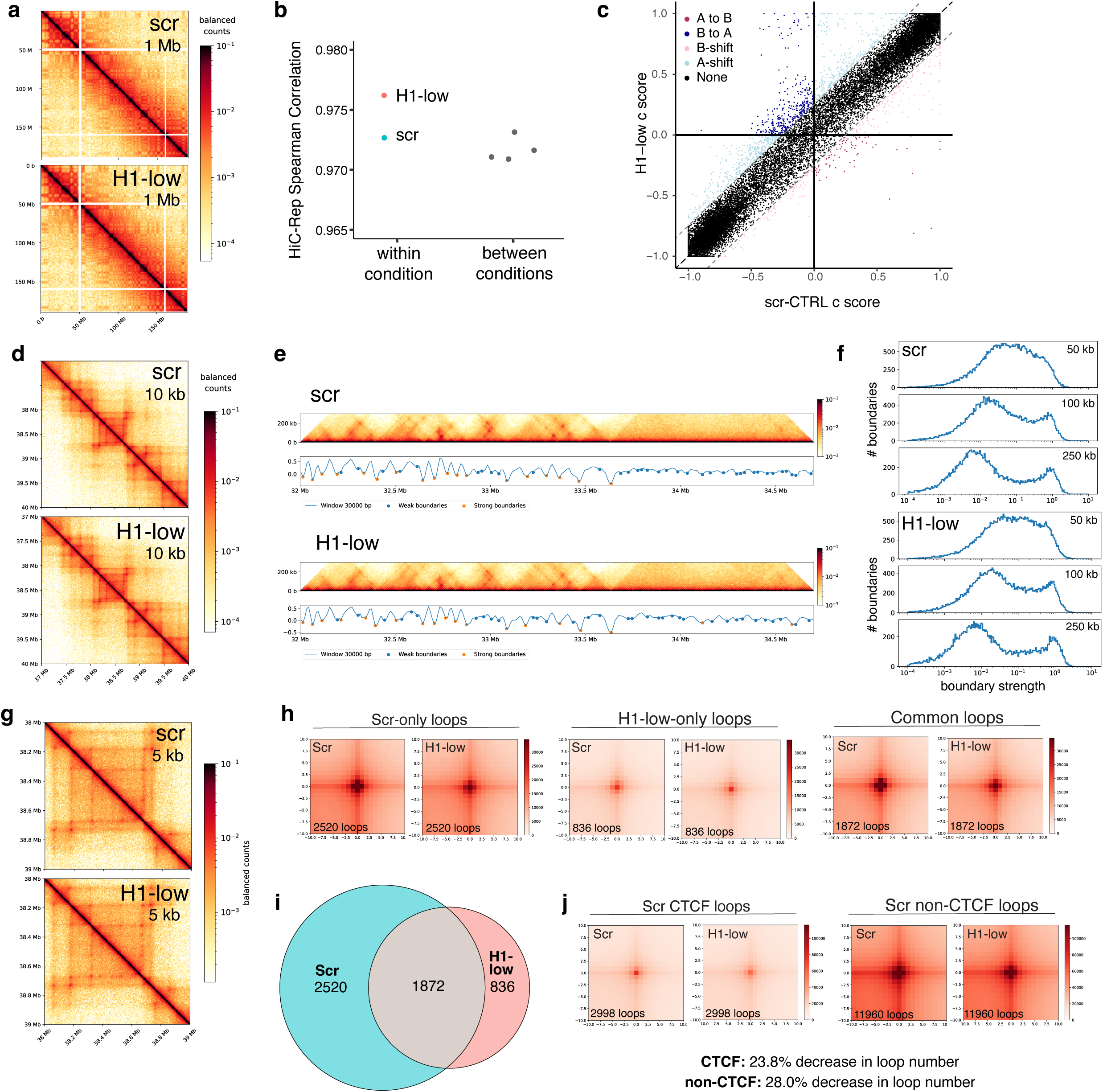
Long-range chromatin structure is only weakly affected by 5-day H1 depletion. **a)** HiC contact maps of scr-CTRL and H1-low cells, ICE corrected by cooltools. Chr 4, 1 Mb resolution, 80-100 M paired end reads per HiC replicate. **b)** Genome-wide reproducibility analysis within (teal and pink) and between (gray) conditions, using HiC-Rep at 500 kb resolution. **c)** Compartment scores (c-scores) in matched genomic bins between scr-CTRL and H1-low HiC contact data at 100 kb resolution. C-score is calculated with cscoretool and compartment shifts are defined as |Δc-score| > 0.2, compartment changes where a bin’s c-score changes sign and A-shift or B-shift where bins shift within the same compartment. **d)** Example contact maps showing domains in scr-CTRL and H1-low Micro-C. Chr 4 zoom, 100 kb resolution, 80-100 M paired end reads per HiC replicate **e)** Example domain boundaries called by cooltools in scr-CTRL and H1-low Micro-C. Chr 1: 32 Mb – 34.7 Mb. **f)** Genome-wide boundary strength distributions for different window sizes for scr-CTRL and H1-low Micro-C. **g)** Example contact maps showing loops in scr-CTRL and H1-low Micro-C. Chr 4 zoom, 10 kb resolution, 80-100 M paired end reads per HiC replicate. **h)** Aggregate peak analysis of loops called by HICCUPS centered around loops called only in scr-CTRL, only in H1-low, or in both datasets. **i)** Overlap of loops called by juicertools HICCUPS in scramble control and H1-low Micro-C. Overlap is defined as both anchors having 90% overlap. **j)** Aggregate peak analysis of loops called by HICCUPS centered around CTCF loops, defined as those overlapping RAD21 and SMC ChIP peaks (called by juicertools motif), and non-CTCF loops. APA is site +/- 10 kb with 1 kb bins.

We then analyzed compartment changes between scr-CTRL and H1-low using the compartment score (c-score). We found that, in contrast to what was observed using the same analysis for H1 depletion by conditional triple knockout in mouse T-cells^6^, the changes in c-score in K562 cells depleted of H1 by five days of CRISPRi were subtle, and only weakly weighted toward B-to-A transitions (Figure 5c).

Visual analysis of chromatin domains shows little change with H1 depletion (Figure 5d), and calls of domain boundaries location (Figure 5e) and strength (Figure 5f) showed that there are no substantial domain changes on the global scale. Calling chromatin loops showed a general loss of loops with H1 depletion (Figure 5i), though the low specificity of loop calling suggests this may in fact reflect an overall weaking of loop strength (Figure 5h). This small loss of loop strength affects both CTCF and non-CTCF loops(Figure 5j).

### Transcriptional de-repression upon H1 depletion preferentially occurs in polycomb repressive complex target genes

Considering the relatively subtle changes in long-range genome organization, we next wondered about the effects of global de-compaction of chromatin and the loss of compaction contrast between epigenetic states on functional outcomes like transcriptional regulation, and its associated features such as DNA accessibility and histone modifications.

We performed poly(A)-capture RNA-seq on scr-CTRL and H1-low cells at five days of H1 depletion to compare against our chromatin compaction results. We found that that the vast majority of changing genes were up-regulated in their transcription (1525 significantly upregulated and 32 downregulated with p < 0.05 and |log_2_(fold-change)| > 1) (Figure 6a). To determine which regulators may be responsible for the changes in gene expression, we performed ChEP-MS to identify changes in protein abundance on chromatin (Figure 6b). We found that the transcription factor GATA1, which is highly expressed in K562 cells, dramatically increased its association with chromatin in H1-low cells, while the BAF complex component SMARCC2, the chromatin-binding nucleoporin NUP153, the H3K4-targeting histone demethylase KDM1B and its methyltransferase KMT2A, the neuron-specific transcription factor TBR1, the polycomb repressive complex2 (PRC2) member SUZ12, and the repressive CBX1 (HP1-beta) protein were decreased in their association. We next used ENRICHR to determine the upstream regulators most likely to explain the change in gene expression (Figure 6c) and followed up with top hits GSEA analyses (Figure 6d). We found that the most significantly upregulated gene sets upon H1 depletion are those regulated by the PRC2 complex member SUZ12 and PRC1 complex members CBX8 and CBX2, which are respectively involved in depositing and sensing the H3K27me3 histone mark. Surprisingly, although the association of GATA1 with chromatin was highly significant, GATA1 targets were not enriched in the upregulated gene set (Figure 6c,e).

**Figure 6:**
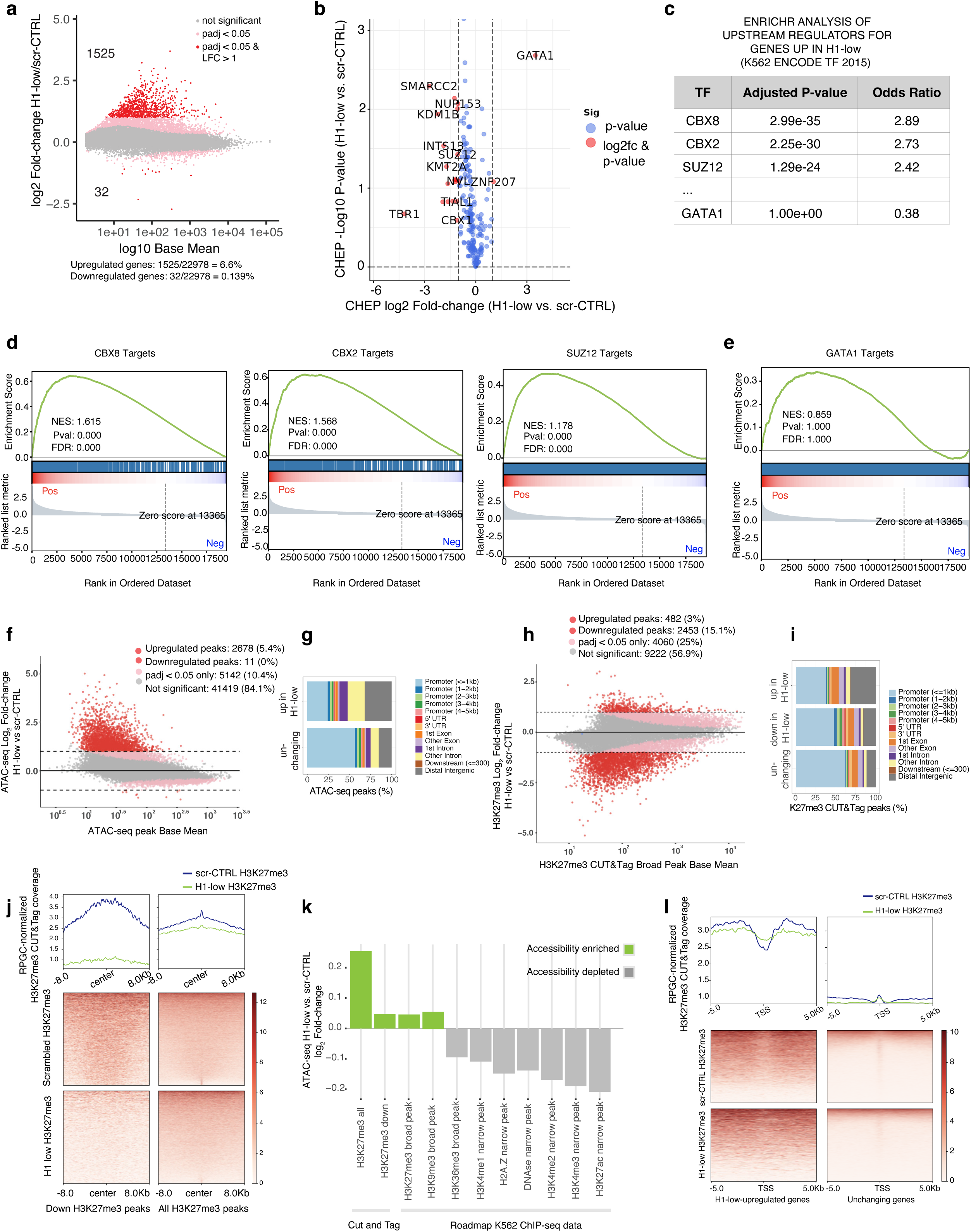
Transcriptional upregulation and increase in accessible chromatin noted upon H1 depletion while opportunistic TF binding does not drive expression changes. **a)** ERCC spike-in normalized RNA-seq. 1525 genes are significantly upregulated upon H1 depletion with |log_2_(fold-change)| >1 and p_adj_ < 0.05 (Benjamini-Hochberg). H1-5, H1-2, H1-3 among the significantly downregulated genes at –1.64, -1.69 and –1.39 log_2_ fold change respectively. **b)** CHEP-seq volcano plot of H1-low vs. scr-CTRL proteins associated with chromatin, n=3 biological replicates for each condition, significance threshold defined as |log_2_(fold-change)| >1. **c)** ENRICHR analysis of transcription factors (TFs) with significantly enriched targets in upregulated genes in H1-low vs. scr-CTRL. TF target gene sets based on ENCODE TF 2015 ChIP-seq peak overlap with target TSS in K562 cells. **d)** Gene set Enrichment analysis (GSEA) of top hits identified by ENRICHR (c) based on Harmonizome ENCODE Transcription Factor Targets database. Normalized enrichment score (NES): CBX8 1.615, CBX2 1.56 and SUZ12 1.17, respectively. **e)** GSEA of GATA1 (identified by CHEP-seq, (b)) target genes based on H1-low vs. scr-CTRL fold-change based on Harmonizome ENCODE Transcription Factor Targets database. NES:0.859 p=1.00. **f)** ATAC-seq peaks base mean over fold change with 2678 peaks significantly upregulated in H1-low condition. |log_2_(fold-change)| >1 and p_adj_. <0.05 (Benjamini-Hochberg). **g)** ATAC-seq peaks annotated by genomic features. **h)** M-A plot of H3K27me3 CUT&Tag fold-change at H3K27me3 CUT&Tag peaks. 2453 peaks significantly downregulated in H1 low condition (|log_2_(fold-change)|>1 and p_adj_. <0.05, Benjamini-Hochberg criterion). **i)** H3K27me3 H1-low upregulated, H1-low downregulated and unchanging peaks annotated by genomic feature. **j)** H3K27me3 signal over H1-low downregulated and all H3K27me3 peaks. Peak centers shown +/- 8kb with a bin size 100 bp. **k)** Enrichment of H1-low/scr-CTRL Log2FC ATAC-seq signal over H3K27me3 CUT&Tag peaks compared to published Roadmap Epigenomics ChIP-seq peaks for K562 ^40^ **l)** H3K27me3 signal over upregulated vs unchanging genes measured by RNA-seq. Signal plot over +/-5 kb around TSS of genes with a bin size of 100 bp.

Next, we investigated how chromatin accessibility responds to loss of H1 using ATAC-seq at 5 days of CRISPRi. We found that as with transcription, differential accessibility is biased toward gains across thousands of ATAC-seq peaks, which are more enriched in distal intergenic regions relative to peaks with stable accessibility (Figure 6f,g).

The increase in accessibility and the de-repression of PRC1/2 target genes led us to hypothesize that loss of H3K27me3 upon H1 depletion may explain the observed increase in transcription. CUT&Tag for H3K27me3 showed widespread changes, with regions changing in both directions but dominated by a loss of H3K27me3 (Figure 6h,i). To tie accessibility changes to epigenetic state, we then asked where the newly accessible sites fell, relative to the existing epigenetic context. We then quantified changes in accessibility in regions that lost H3K27me3 compared to those where the signal remained unchanged, as well as in other regions marked by several additional epigenetic marks (Roadmap Epigenomics ^40^) (Figure 6k). Consistent with the gene regulation results, we saw that the regions with net increases in accessibility are heterochromatic—those marked by H3K27me3 in scr-CTRL or parental K562 cells and those marked by H3K9me3 in parental K562 cells ^40^ (Figure 6k). However, the fold-change of accessibility in regions losing H3K27me3 was not higher when compared to all H3K27me3 regions. Similarly, when we investigated the change in H2K27me3 between control and H1-low cells specifically focusing on genes that increased in transcription, we found that the local H3K27me3 landscape stayed at a similar level (Figure 6l). What was notable, was that the genes that were upregulated upon H1 loss had a much higher level of H3K27me3 signal near their promoters than genes that were not de-repressed, regardless of H1 depletion (Figure 6l).

Together, these results suggest that gene de-repression and the gain of accessibility does not require complete local loss of promoter proximal H3K27me3 and that the mechanism of de-repression is not simply a direct consequence of local H3K27me3 loss.

### Chromatin de-compaction by H1 depletion is genome-wide, except for regions that were already de-compacted and accessible

We next looked at the regions that gain accessibility in H1-low versus scr-CTRL–spanning promoter proximal and distal sites. We found that H3K27me3 signal flanking these peaks of accessibility is largely maintained in H1-low (Figure 7a), indicating that accessibility gains do not generally require local depletion of H3K27me3 at these regulatory elements. The local difference in the CUT&Tag signal observed is likely to be driven by the change in accessibility, as a sharp CUT&Tag peak at the center of the newly opened ATAC-seq peaks (Figure 7a).

**Figure 7:**
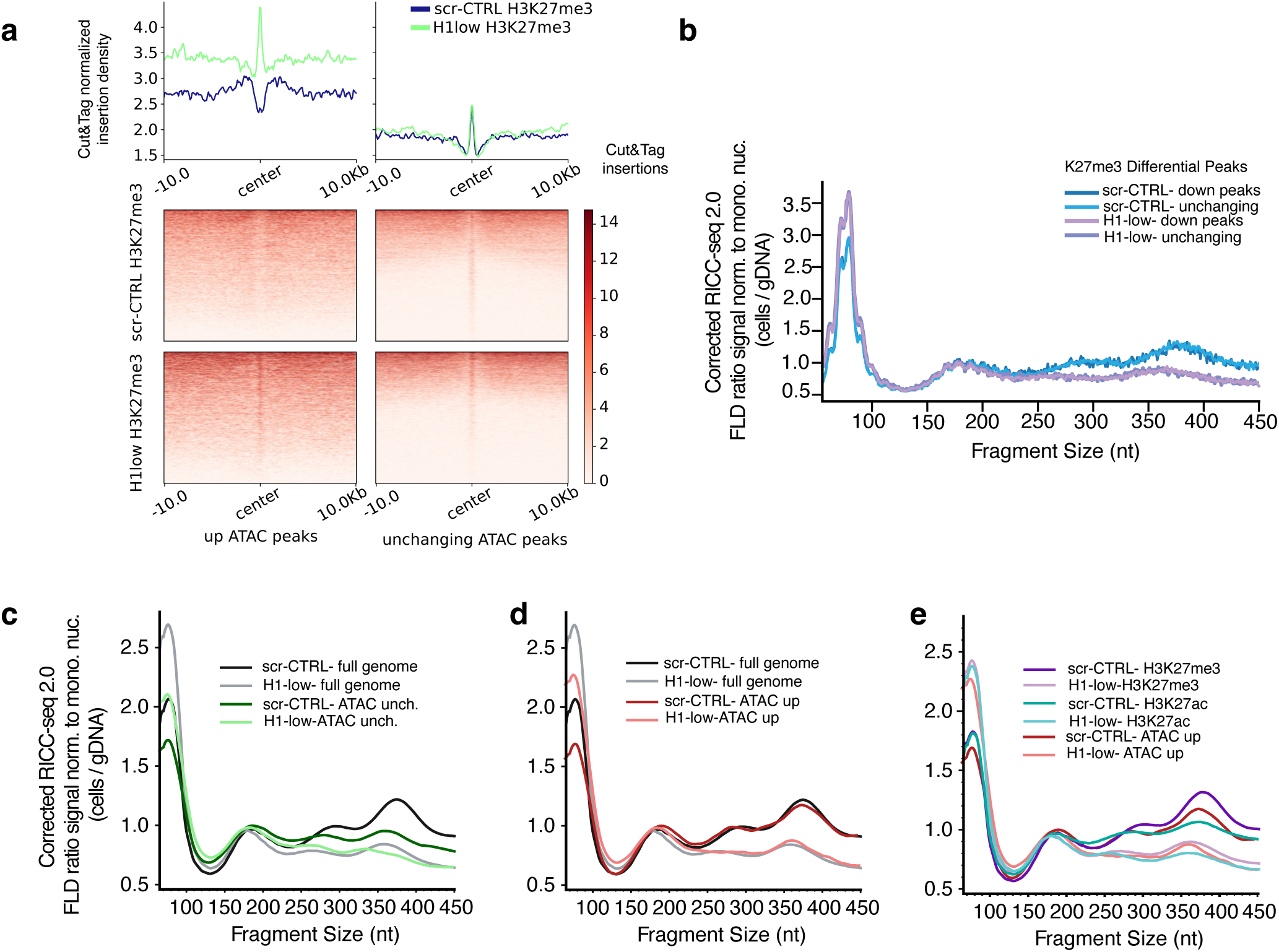
Chromatin fiber compaction is unaffected by H3K27me3 depletion but varies with changes in accessibility. **a)** ATAC-seq accessibility signal over Cut&Tag H3K27me3 peaks +/- 5 kb around peak centers shown with a bin size of 100 bp. **b)** RICC-seq FLD plot over downregulated vs unchanging H3K27me3 peaks in H1-low vs scr-CTRL. n= 1 biological replicate shown as ratio over subset genomic DNA control. Signal normalized to mononucleosome peak. **c)** RICC-seq FLD plot over unchanging ATAC-seq peaks and genome-wide smoothed (30 nt rolling average). n=1 biological replicate shown corrected by biological replicate-specific correction factor and ratio over similarly subset and corrected sample-matched genomic DNA control. Signal normalized to mononucleosome peak. **d)** RICC-seq FLD plot over upregulated ATAC-seq peaks and genome-wide, as in (c). **e)** RICC-seq FLD plot over upregulated ATAC-seq peaks, H3K27me3, and H3K27ac, as in (c).

We then turned to RICC-seq as a measure of chromatin compaction to determine whether there are focal changes in chromatin compaction at regions where DNA accessibility or H3K27 trimethylation are changing. We did not observe any change in chromatin compaction by RICC-seq between genomic regions that lose K27me3 and those that do not (Figure 7b). The primary difference remains between the scr-CTRL and the H1-low sample at all H3K27me3-marked sites, regardless of their change in the histone mark between the two conditions.

For genomic regions that change accessibility, on the other hand, we did observe changes in compaction (Figure 7c-e). We found a distinction in the average RICC-seq FLD between unchanging ATAC-seq peaks (Figure 7c) and those that become accessible upon H1 depletion (Figure 7d). Unchanging ATAC-seq peaks are already quite decompacted, with FLDs similar to genome-wide acetylated chromatin (Figure 7e) and resembling the decompacted chromatin of budding yeast (Figure 2). The regions that gain accessibility, however, began with a FLD very similar to the genome-wide average and decompacted to a FLD comparable to the genome-wide H1-low FLD upon H1 depletion (Figure 7e).

## Discussion

We set out to understand the relationship between chromatin structure, transcriptional regulation, DNA accessibility and histone marks. By improving the RICC-seq protocol to RICC-seq 2.0, we obtained a protocol that could be reliably applied to a variety of sample types with varying levels of chromatin compaction. We verified that cell types with very different NRLs and global levels of chromatin compaction, spanning de-compacted budding yeast cells through mammalian cell types and hyper-compacted, transcriptionally inactive chicken erythrocytes, produced different RICC-seq FLDs, demonstrating that the method is sensitive to changes in chromatin compaction.

We were particularly motivated to make direct *in situ* measurements of chromatin compaction in a model of linker histone depletion because linker histone H1 has so often been invoked as an architectural protein that uses chromatin compaction as its mechanism for broad-based transcriptional repression. Our results show that a dramatic reduction in total H1 levels leads to not only an increase in accessibility at thousands of sites and the upregulation of thousands of genes, but it also causes chromatin decompaction at the tri-nucleosome length scale, which we could measure using two orthogonal methods—Micro-C and RICC-seq 2.0. We did not see strong changes in long-range chromatin organization over the same time scale, suggesting that that chromatin structure at this scale is not directly coupled to H1 density and transcriptional regulation. This underscores the importance of maintenance of chromatin compaction by linker histone in regulating both the accessibility of many sites across the genome and the transcriptional repression of a large set of genes. We also observed a modest but broad-based loss in H3K27me3 signal as measured by CUT&Tag, indicating that in this system, linker histone plays a role in the maintenance of the H3K27me3 mark, as has been observed in other systems, including T-cells^6^ and B-cells^41^, as well as in K562 cells in which H1 is depleted via CRAMP1 knockout^42^.

Surprisingly, the changes in chromatin compaction upon linker histone depletion are remarkably uniform across the genome. Although DNA accessibility is particularly enriched in H3K27me3-decorated regions of the genome upon H1 depletion, this is not accompanied by specific decompaction at H3K27me3 regions, any more than is happening in the rest of the genome. RICC-seq is, however, sensitive to changes in compaction elsewhere. We observed a difference in the FLD shift between regions that maintained accessibility and those that gained it—those with pre-existing accessibility had a more de-compacted FLD in control cells, and experienced only a modest change in the FLD and in the compaction contact peak upon H1 depletion.

Our results support a model in which linker histone H1 is not locally inducing compaction at particular loci, but is rather working genome-wide to compact most chromatin, with the exception of acetylated regions where it is removed and chromatin can de-compact^10,39^. This is consistent with FRAP data showing that *in vivo*, linker histones are remarkably dynamic^43^, which would permit their broad distribution across the genome, and with electron microscopy and super-resolution microscopy, which show a broad change in both chromatin density and nucleosome clutch size ^6,21^. There may be variability in H1 density between chromatin types, and chromatin compaction at the tri-nucleosome scale may have different sensitivity to the linker histone level as compared to other architectural or functional features. Indeed, some threshold effects in chromatin fiber architecture have been observed with linker histone density changes *in silico*^44^.

Overall, we find that H1 acts to modulate the global nucleosome repeat length and local compaction of chromatin, but that some chromatin states may be more dependent on this compaction than others. PRC1/2 repression of both accessibility and gene expression is particularly sensitive. The exact nature of this sensitivity may be a combination of H1’s effects on both chromatin compaction and NRL. *In vitro* experiments show that PRC2 deposition of H3K27me3 preferentially occurs on long-NRL chromatin^45^, but compaction also promotes deposition of H3K27me3 and prevents deposition of its antagonistic mark H3K36me2 ^6^.

Our results highlight that there is a close regulatory relationship between H1-dependent sub-kilobase chromatin compaction, DNA accessibility, histone marks and transcriptional regulation—and that it is more immediate than the long-range compartmentalization of the nucleus. In a system with as much complexity and redundancy as chromatin, directly measuring chromatin compaction as a distinct physical variable, rather than inferring it from other methods, can help define more precise mechanistic models of transcriptional repression.

## Acknowledgments

We would like to thank the Risca Lab, and Skoultchi Lab, and the members of the Rockefeller Chromatin Supergroup, as well as Ari Melnick, Ethel Cesarman, and Yael David for helpful discussions. We also thank the support of the Rockefeller University Genomics Resource Center and High Performance Computing Resource Center. This work was supported by a NIH New Innovator Award to V.I.R. (DP2GM150021), a Rita Allen Foundation Scholar Award to V.I.R., a Hirschl/Weill-Caulier Career Scientist Award to V.I.R., a NSERC post graduate scholarship award to H.C., and a HFSP Postdoctoral Fellowship to A.O. H.D. P. and A.I.S. were supported by NIH grant R01GM147165 and D.V.F. by NIH grant R01HD114814.

## Methods

### Cell culture and preparation

#### BJ-5ta Fibroblasts

Cells were grown in DMEM supplemented with 10% FBS and 1% Penicillin-Streptomycin. Cells passaged every three days at 1:3 splits when cells are about 75% confluent. To harvest, cells were contact inhibited and trypsinized with 4 ml Trypsin to lift then quench with 8 mL media. Cells were washed and spun down with PBS and 2 million cells per technical replicate harvested per plug.

#### Budding yeast

For RICC-seq experiments on budding yeast, we used *Saccharomyces cerevisiae* W303 *RAD5+* wild-type strains (Gift from Xiaolan Zhao). Overnight starter cultures were diluted into fresh YPD to an initial OD_600_ of ∼0.1 and grown at 30°C with shaking for at least 4 hours to allow cells to enter mid-log growth phase with an OD_600_ of 0.5–1.0. Cells were harvested in 50 mL conical tubes by centrifugation at ∼900 × g for 3 min, washed once in 25 mL PBS, transferred to 1.5 mL microcentrifuge tubes, and washed again in 1 mL PBS (30 s at ∼900 × g between washes). Pellets were resuspended in PBS and mixed 1:1 with molten low-melting-point (LMP) agarose. Yeast cells were embedded directly in agarose plugs without prior zymolyase treatment, as pilot experiments indicated it was not required for efficient lysis and downstream processing. The cell–agarose suspensions were immediately cast into plug molds and solidified on ice for 10 min. Plugs were released into 2 mL tubes (3 plugs per tube) and irradiated on ice in a 50 mL conical tube with 1000 Gy X-ray ionizing radiation at ∼120 Gy/min over the course of 8 minutes and 20 seconds, while non-irradiated tubes were kept on ice for the same duration. Following irradiation, plugs were immediately incubated in 950 µL RICC lysis buffer supplemented with 50 µL Proteinase K at 25°C with gentle shaking for 48 h. After cell lysis, the rest of the RICC-seq protocol proceeded as described further below.

#### Chicken Red blood cell

Whole chicken blood was ordered from Pel-Freez Biologicals, Whole Chicken Blood, Non-Sterile with Alsever’s Media, Cat. No. 33133-1. Cell concentration was determined using a hemocytometer corresponding to ∼9.8 × 10^8 cells/mL. A 5 mL aliquot was transferred to a 15 mL conical tube and pelleted at 200 × g for 5 min, the supernatant was removed, and the cells were washed twice in PBS at 200 × g, 5 min for each wash. The pellet was spun again at 200 × g for 5 min and kept on ice, then resuspended in 2.5 mL PBS to generate a suspension at 1.5 × 10^2 cells/mL. Cells were aliquoted at 750 µL into 2 mL tubes, equilibrated at 37°C for 1 min, and mixed 1:1 (v/v) with pre-warmed 2% low–melting point agarose. The solution was pipetted carefully into plug molds, avoiding bubbles and allowed to solidify on ice. Plugs were then transferred into 400 µL cold PBS and either irradiated with 300 Gy ionizing radiation on ice or kept on ice as non-irradiated 0 Gy and PLC controls. Following irradiation, plugs were incubated in 1,170 µL lysis buffer supplemented with 30 µL Proteinase K; samples were kept on ice for 2–3 h post-lysis and then transferred to room temperature for overnight incubation.

#### K562 H1 depletion and scrambled control cell culture and induction

K562 cells expressing a dox-inducible dCas9-KRAB-P2A-mCherry were generated by lentiviral transduction with the TET-ON vector pAAVS1-NDi-CRISPRi (addgene #73497). Transduced cells were selected with 200ug/ml G418 and inducible dCas9-KRAB-P2A-mCherry cells were further selected with fluorescence-activated cell sorting (FACS) after 3 days of Doxycycline treatment (1ug/ml). Selected cells were allowed to grow in the absence of doxycycline and then transduced with pU6-sgRNA EF1Alpha-puro-T2A-BFP (addgene #60955) which was engineered with 4 in tandem U6-sgRNA expression cassetes, each expressing a subtype specific H1 sgRNA namely H1.5 (GGCAGGAGCGGTTTCCGACA), H1.2 (GGCTGCCGCCGGCTATGATG), H1.3 (GGCTGCCGCCGGCTATGATG) and H1.4 (GGCCAAGCCTAAGGCTAAAA). As control, cells were also transduced with a non targeting pU6-sgRNA EF1Alpha-puro-T2A-BFP (GCACTACCAGAGCTAACTCA). Cells expressing constitutive high levels of sgRNAs were selected by combining puromycin selection (10ug/ml) and FACS to collect the top BFP positive cells. H1 depletion was induced by supplementing culture media with doxycycline 1ug/ml for 5 days and H1 content assayed by RP-HPLC of acid extracted histones.

Cells were grown in IMDM media (Thermo Scientific #12440046) supplemented with 10% heat-inactivated FBS (Sigma-Aldrich F4135) and 1% Penicillin-Streptomycin. Cells were passaged every two days, seeding 2.4 M cells in a T150 flask. Starting five days before experimental harvest, doxycycline was added to the growth media of both H1-low and scr-CTRL cells to a concentration of 3 mg/mL. Cells were grown in the doxycycline media for five days, following the regular splitting schedule, then harvested for experiments on day five. 2 million cells per technical replicate harvested.

#### Naïve B cell isolation

Spleens from wild-type C57BL/6J mice (Jackson Laboratories, strain 000664) were mechanically dissociated and passed through a 40-µm strainer. Red blood cells were lysed using ACK buffer (Lonza). Resting B cells were then enriched by negative selection using anti-CD43 (Ly48) magnetic microbeads (MACS, Miltenyi Biotech), according to the manufacturer’s instructions. Briefly, the cell suspension was incubated with 30 µl of CD43 magnetic beads diluted in 270 µl of PBS per spleen for 20 mins at 4 °C. The mixture was then applied to an LS MACS Column on MACS Separator. The flow-through containing naïve B cells was collected and resuspended in PBS supplemented with 0.5% bovine serum albumin (BSA) and 2 mM EDTA.

### RICC-seq 2.0 library preparation

RICC-seq 2.0 was performed on multiple different cell types each with their own growing techniques and harvest conditions as outlined above. Adherent cells were grown to ∼100% confluence and rested ∼1 day to enrich G0. Cells were were washed in warm PBS (Gibco 14190-144), detached with 0.05% Trypsin-EDTA or Accutase at 37 °C (Invitrogen 25300-054), counted with trypan blue (Gibco 15-250-061), pooled, and pelleted (200–300 RCF, 4 min). Pellets were resuspended in PBS to 50 M cells/mL and embedded to final concentration 1% low-melt agarose (Sigma Type VII-A, A0701-25G) kept at 37 °C, using Bio-Rad plug molds (1703713). 2 million cells per plug were used for the K562 cells harvested 5 days post doxycycline induction.

Plugs were irradiated to a total dose of 300 Gy for most conditions except 1000 Gy for yeast and lysed overnight (24-48 h, RT/20 °C with shaking) in RICC lysis buffer containing 20% N-lauroylsarcosine (Sigma L7414-50ML), Proteinase K (NEB P8107), and EDTA (Thermo 15575020), then washed for ∼5 h as follows: TE + 1 mM PMSF, 30 min at 4 °C; TE + 1 mM PMSF, 45 min at 4 °C; TE, 60 min RT; TE, 60 min RT; TE + RNase A 0.1 mg/mL (Sigma R4642-250MG), 45 min at 37 °C; TE + RNase A 0.1 mg/mL, 45 min at 37 °C; TE, 60 min RT.

Genomic DNA control plugs were equilibrated in 0.5 M Tris-HCl pH 8 on ice (three 15-min exchanges, then +400 µL Tris) and irradiated identically, followed by the same washes. ssDNA was eluted by incubating plugs in 0.1 N NaOH (200 µL, 15 min), neutralized with 1 M Tris-HCl pH 7.5 (100 µL) + 2 mM EDTA (RT, ∼4 h), and purified with the Zymo RNA Clean & Concentrator-5 kit (R1016) using modified speeds (binds at 3,800 RCF; washes at 10,000 RCF); columns were loaded with 600 µL RNA Binding Buffer then 900 µL absolute ethanol and eluted twice in 10 µL 10 mM Tris pH 8 (total ∼18 µL). K562 eluate samples were purified with custom-made carboxyl-coated beads to facilitate batch processing^46^.

Spike-ins were included post elution into each sample before library preparation. A genomic yeast DNA ladder digested with MNase was included for cross-experiment comparability.

Ends were dephosphorylated with rSAP (NEB M0371) in CutSmart buffer (21 µL total; 37 °C, 1 h), then 5′-phosphorylated with T4 PNK (Enzymatics Y9040L) in the presence of DTT (125 mM stock to 5 mM final; Sigma 43815-1G) and ATP (500 µM stock to 5 µM final) in a 25 µL reaction (37 °C, 1 h); MgCl₂ (Invitrogen AM9530G) and spermidine (Sigma 85558-1G).

SRSLY ligation was done with SRSLY P5/P7 adapters, T4 DNA ligase (Enzymatics L6030-LC-L) with 1× ligase buffer and 18.5% PEG-8000 (50% stock) at 37 °C for 1 h; adapters were added at ∼250 nM each. Post-ligation clean-up used Zymo R1016 as above, eluting in 12 µL 10 mM Tris pH 8. K562 were cleaned up using custom-made size selection beads^46^.

Libraries were barcoded by PCR in 50 µL with KAPA HiFi PCR Mix (2×; KAPA KK2602), 2 µM each i5/i7 barcode primer, and 21 µL template; cycling was 98 °C 3 min; 5 initial cycles of 98 °C 30 s, 65 °C 30 s, 72 °C 1 min; 72 °C 1 min. A side-reaction qPCR on a 5 µL aliquot determined additional cycles to retain exponential amplification (BJ, K562 typically 12 total cycles). gDNA control, 0 Gy, and irradiated conditions were processed in parallel through all steps. Rad source 1800 Q4 X-ray Irradiator used delivering a dose rate of 123.1 Gy/min on the top shelf. BJ, K562, Chicken red blood cells were irradiated with a total dose of 300 Gy and Budding Yeast at 1000 Gy.

### Hi-C library preparation

HiC was performed using the HiC 3.0 protocol followed by NEBNext Ultra II library preparation. Briefly, pellets of 5 M cells were crosslinked first with 1% formaldehyde in HBSS for 10 minutes, then with 3 mM DSG for 40 minutes, and subsequently snap frozen in liquid nitrogen. Cells were lysed with a Dounce homogenizer in a buffer containing 0.2% NP-40, followed by chromatin solubilization by SDS, quenched with Triton X-100. Chromatin was digested with a cocktail of DdeI (400 U) and DpnII (400 U), incubated overnight at 37°C. DNA ends were then biotinylated using Klenow polymerase and a dNTP mix containing biotin-14-dATP, then proximity ligation was performed using T4 DNA ligase. After biotin fill-in and proximity ligation, crosslinks were reversed and proteins digested with an overnight 65°C incubation with Proteinase K. DNA was then purified and concentrated via phenol-chloroform extraction followed by ethanol precipitation and passage through Amicon filter tubes. End repair was performed with T4 polymerase and dATP/dGTP, then DNA was sonicated to obtain a distribution of sequenceable DNA lengths, which was further size selected using AMPure XP beads. At this point, we switched to NEBNext Ultra II library preparation, following the manufacturer’s protocol, then sequenced samples on an Illumina NovaSeq X Plus platform to a depth of 50 - 90 million read pairs per replicate for two technical replicates.

### In-situ Micro-C library preparation

Two biological replicates of Micro-C were performed using an in-situ proximity ligation-based protocol, adapted for our purposes from previously published Micro-C protocols, as described below.

#### Crosslinking

Pellets of 10 million cells were crosslinked with 1% formaldehyde for 10 minutes, quenched with 1 M Tris-HCl pH 7.5, washed with DPBS, then crosslinked with 3 mM DSG (synthesized in house) for 45 minutes. DSG crosslinking was quenched with 1 M Tris-HCl pH 7.5, and cells were washed with DPBS before snap freezing in liquid nitrogen.

#### MNase titration

Cell pellets were thawed on ice and resuspended in buffer MB1^28^. The cell suspension was split into five aliquots of 2 million nominal cells, 2-3 of which aliquots were used for MNase titration, and 2-3 of which were used as experimental samples. Experimental samples were kept on ice while the MNase titration was took place (typically about three hours). Titration was carried out by extracting nuclei by incubating cells in MB1 for 20 minutes, spinning down and resuspending nuclei in MB1, then adding varying volumes of MNase (NEB M0247S) to aliquots. Typically, volumes between 2 and 6 uL of 1:10 diluted MNase produced the desired level of digestion. MNase digestion was performed for 10 minutes at 37°C on a thermomixer, then stopped with EGTA and incubation at 65°C. Digested chromatin was treated with Proteinase K (NEB P8107) and RNase A (Sigma-Aldrich R4642) and incubated for 2 hours at 65°C to reverse crosslinks and digest proteins. DNA was then purified with phenol-chloroform extraction followed by passage through a Zymo DNA Clean & Concentrator kit. The digestion profiles of the titration samples were assessed by quantifying the DNA by Qubit and running 1 ng on a TapeStation. MNase concentrations leading to a major mononucleosome peak and small dinucleosome peak were identified and used for the full protocol with the remaining aliquots left on ice.

#### Micro-C

Nuclei were spun down, resuspended in MB1 with the predetermined concentration of MNase, and digested as in the titration. The digested sample was washed with MB2^28^ then end repaired and end labeled with PNK (NEB M0201L) and Klenow polymerase (NEB M0210M) with biotinylated dATP and dCTP. After enzyme inactivation with EDTA and incubation at 65°C, the sample was washed with MB3^28^. Proximity ligation was carried out using T4 ligase (NEB M0202L) and unligated fragments were digested with Exonuclease III (NEB M0206L). The sample was then incubated overnight with Proteinase K and RNase A at 65°C to reverse crosslinks and digest proteins. DNA was purified with phenol-chloroform extraction followed by passage through a Zymo DNA Clean & Concentrator kit, then ligated mononucleosome pairs were selected for by running the sample on a 2% agarose gel, excising and extracting the dinucleosome-sized band, and pulling down with streptavidin beads. Library preparation was then carried out using the NEBNext Ultra II kit, following manufacturer’s instructions. Libraries were sequenced twice on an Illumina Novaseq X Plus to approximately 100-400 million 2 x 150 bp paired end reads per replicate per run. Reads from the two runs were combined for a final depth of 500 - 600 million reads per technical replicate, or 1.6 - 2 billion reads per biological replicate.

### ATAC-seq library preparation

ATAC-seq was performed following the Omni-ATAC-seq protocol, using Tn5 made in-house with the Open-Tn5 method. Tn5 was loaded with Illumina adaptors by incubating equal volumes of 1 mg/mL Tn5 and 1 𝜇M adaptor together for 10 minutes at room temperature immediately before being used for tagmentation. Briefly, the Omni-ATAC-seq protocol involved harvesting cells in aliquots of 50,000 cells – in this case, we did two separate harvests (biological replicates) with 3-5 aliquots (technical replicates) per condition – followed by light permeabilization in a buffer containing 0.1% NP-40, 0.1% Tween, and 0.01% digitonin. The samples were then tagmented with Tn5 at 37°C for 30 minutes, cleaned up with a Zymo DNA Clean & Concentrate kit, and PCR amplified with barcoded primers for an optimized number of cycles determined with a side qPCR reaction. Following amplification, libraries were cleaned up, quantified, and sequenced on an Illumina NextSeq 2000 platform to a depth of 30 - 50 million read pairs per technical replicate.

### CUT&Tag library preparation

Cut&Tag libraries were prepared using the Epicypher protocol. K562 cells were lightly fixed using 0.1% formaldehyde for 1 minute, spun down resuspended in cell freezing media and frozen down post day 5 doxycycline induction of H1 depletion. Pellets were then hawed, spun down, washed in PBS and 100,000 cells per technical replicate counted. The cut and tag protocol proceeded as outlined in the Epicypher v1.7 protocol. TN5 for these CUT&Tag libraries was made in house using our previously published protocol^47^. The antibody used for mapping H3K27me3 was CST 9733 at a 1:100 final concentration.

### RNA extraction and RNA-seq library preparation

Triplicate cultures of cells expressing H1 sgRNAs and Scramble sgRNA were induced with Doxycycline 1ug/ml for 5 days. Cells were backdiluted during treatment in order to maintain cell cultures in exponential growth phase. Five mmillion cells were collected on day 5. One million cells were ressuspended in Trizol for RNA extraction and four million cells used for acid extraction and HPLC for validation of H1 depletion. Total RNA was extracted using Direct-zol RNA Miniprep Kit (Zymo Research). Standard mRNA-Seq (poly(A) selection) was performed at Azenta Life Sciences. Libraries were performed incorporating unique molecular identifers during adapter ligation and External RNA Controls Consortium (ERCC) spike-ins were added to ech sample before reverse transcription.

### Chromatin Enrichment for Proteomics (ChEP)

Chromatin-bound proteins were isolated using the ChEP protocol as described by Kustatscher et al.^48^ with minor modifications. Briefly, cells were formaldehyde-crosslinked (1%, 10 min), nuclei were isolated under hypotonic conditions, and chromatin was enriched by sequential high-stringency washes under denaturing conditions (2% SDS, 8 M urea). Crosslinked chromatin was sonicated and used for quantitative Mass Spectrometry analysis.

### Mass-spectrometry

Samples were alkylated with 30mM IAA for 45min at RT in the dark. Reactions were then desalted into 50mM NH4HCO3 using ZebaSpin 7k columns (ThermoFisher) and eluates were supplemented with trypsin (0.1mg/ml) and digested for 2h at 37C. At the end of the 2h, samples were supplemented with additional trypsin and digestions allowed to proceed overnight. Digestions were quenched with 1% formic acid, dried in SpeedVac and then resuspended in 130 µl MS Sample Buffer (0.1% formic acid, 1% acetonitrile in water).

LCMS analyses were performed on a TripleTOF 5600+ mass spectrometer (AB SCIEX) coupled with M5 MicroLC system (AB SCIEX/Eksigent) and PAL3 autosampler. LC separation was performed in a trap-elute configuration, which consists of a trap column (LUNA C18(2), 100 Å, 5 μm, 20 X 0.3 mm cartridge, Phenomenex) and an analytical column (Kinetex 2.6 μm XB-C18, 100 Å, 50 X 0.3 mm microflow coumn, Phenomenex). The mobile phase consisted of water with 0.1% FA (phase A) and 100% ACN containing 0.1% FA (phase B).

Peptides in MS Sample Buffer were injected into a 50-μl sample loop, trapped and cleaned on the trap column with 3% mobile phase B at a flow rate of 25 μl/min for 4 min before being separated on the analytical column with a gradient elution at a flow rate of 5 μl/min. The gradient was set as follows: 0–24 min: 3% to 35% phase B, 24–27 min: 35% to 80% phase B, 27–32 min: 80% phase B, 32–33 min: 80% to 3% phase B, and 33–38 min at 3% phase B. An equal volume of each sample (30 μl) was injected four times, once for information-dependent acquisition (IDA), immediately followed by DIA/SWATH in triplicate. Acquisitions of distinct samples were separated by a blank injection (80 µl MS Sample Buffer) to prevent sample carryover. The mass spectrometer was operated in positive ion mode with EIS voltage at 5200 V, Source Gas 1 at 30 psi, Source Gas 2 at 20 psi, Curtain Gas at 25 psi, and source temperature at 200°C.

### RICC-seq data processing

#### RICC-seq library alignment and fragment length distribution (FLD) generation

##### Alignment

Illumina FASTQ paired end reads were aligned with Bowtie2 to prebuilt Bowtie2Index references (e.g., hg38/hg19/mm10), applying a mapping-quality cutoff (MAPQ ≥ 30) and optional removal of reference blacklist regions (defaults provided per genome when available). Per-sample reads to the yeast genome E2F were also aligned and used for downstream spike in correction and length bias normalization.

##### Subsetting

For each input BAM, we performed a name sort (samtools sort -n) to ensure proper pairing semantics for downstream intersection. Paired-end alignments were then intersected with each provided BED/peak set using bedtools pairtobed (criterion: overlap of the paired fragment span with the feature set; -type ospan). The script iterates over all BAM×BED combinations. The resulting per-combination subset BAMs (reads whose paired fragment spans overlapped the specified regions) were carried forward to downstream analyses.

##### Fragment length histogram generation

For each input BAM, we computed paired-end insert-size distributions with Picard Toolkit (Broad Institute) CollectInsertSizeMetrics. The task excludes PCR/optical duplicates. Picard was invoked with DEVIATIONS=10.0 to capture long-tail fragments up to mean ± 10 SD, MINIMUM_PCT=0.05 to require ≥5% of pairs for a stable estimate, and HISTOGRAM_WIDTH=700. For each BAM, the script writes (i) a PDF histogram <BASENAME>_hist.pdf and (ii) a tabulated metrics log <BASENAME>_full_hist_graphwithoutdups.log (median/mean/stdev, read counts, and percentile cutoffs) for downstream QC. The log files for both spike in controls and samples were then compiled to use for plotting Fragment Length Distributions (FLD) and correcting.

#### FLD normalization and correction

Fragment length distributions were corrected in two stages. First, per-sample spike-in normalization was applied using a biological replicate-specific scaling factor defined as the ratio of the mean spike-in read depth of all technical replicates within a biological replicate (REF_𝑏_) to the individual sample’s spike-in depth (𝐷_𝑖_). Each replicate distribution was multiplied by REF_𝑏_/𝐷_𝑖_ to equalize spike-in coverage across replicates.

Second, to correct for fragment-length bias, we used the spike-in–scaled curve and its length-bias–corrected counterpart calculated by comparing the spike-in ladder FLD fragment loss after sequencing to the ladder input on TapeStation. For each technical replicate, a length-bias correction factor was computed per base pair. Within each biological replicate, these per-technical replicate correction curves were averaged to obtain a mean biological replicate-specific falloff profile which was used to correct each averaged biological replicate curve.

Each curve was then normalized to the signal of the mononucleosome at 180 bp and lightly smoothed (10 bp rolling average). These biological replicate-level normalized profiles were used to compute condition means ± 95 % confidence intervals and to perform per-base Welch’s *t*-tests, with significant contiguous regions (≥ 5 bp) highlighted in downstream figures. To summarize a condition (e.g., SCRM or dH1), we stacked all available replicates from that condition and compute the condition mean curve as the pointwise average across biological replicates. 95% confidence intervals were obtained using a Student’s t interval across biological replicates, i.e., mean ± 𝑡_0.975, 𝑛−1_ ⋅ SD/√𝑛, where 𝑛 is the number of biological replicates at that bp (using exact 𝑡for small 𝑛, ∼1.96 as 𝑛grows). For between-condition comparisons we plotted the pointwise dH1/SCRM mean ratio and assessed significance at each bp with a two-sided Welch’s t-test computed over the underlying biological replicate curves, shading only significant segments of length ≥ 5 bp as significant differences.

All shown subset RICC-seq curves were length bias corrected by the previously calculated respective biological replicate correction factor and divided by the corrected genomic DNA control (PLC) curve to produce the signal over the genomic background per condition.

### HiC and Micro-C analysis

#### Alignment and QC filtering

Basic alignment and quality control (QC) filtering for both HiC and Micro-C was done based on the Dovetail Genomics analysis pipeline (micro-c.readthedocs.io), in which ligation pairs were aligned with bwa mem (v 0.7.17) using two-sided alignment, then valid ligation events were identified with pairtools parse (v 0.3.0). PCR duplicates were removed and final bam and pairs files were created with pairtools split. From pairs files, ICE balanced mcool files with a base resolution of 500 bp were made using cooler cload and zoomify (v 0.8.6) with default parameters, and hic files with a base resolution of 500 bp were made using juicer_tools pre (v 1.22.01) with default parameters. Pairs, mcool, and hic files were then used for subsequent analyses, described below.

#### HiC analysis

P(s) curves were made from balanced mcool files at 10 kb resolution using cooltools (v 0.5.4) expected_cis with smoothing. For correlation analysis with HiCRep^49^ , we converted our HiC pairs files to full contact matrices at 500 kb resolution using juicer and straw (v 1.6)^50^, then computed pairwise SCC scores between all replicates, within and between conditions. Chromosome compartment analysis was carried out using the cscoretool package (v1.1)^51^ . First, cscoretool was run on each chromosome for each condition, using pairs files at 100 kb resolution. We next calculated Spearman’s correlation coefficient for each chromosome’s c-scores with H3K36me3 ChIP signal (from ENCODE^52^ ENCSR000AKR) and flipped the sign of the c-score for negatively correlated chromosomes so that positive c-scores consistently correspond to the gene-dense A compartment (supp fig ref). Chromosomes for which correlation did not pass a significance threshold of p < 0.01 were excluded from the analysis. Differential compartment scores were found by subtracting scrambled control c-scores from H1-low c-scores in matched genomic bins, and shifts were defined as |Δ c-score | > 0.25, and negative to positive = B to A, positive to negative = A to B, increasing within positive or negative = A-shifted, decreasing within positive or negative = B-shifted.

#### Micro-C domain and loop calling

Domain analysis was performed using the cooler/cooltools suites. From balanced mcool files, the cooltools insulation tool was used to find insulation strength and call domain boundaries at a base resolution of 10 kb. Loop calling was performed on hic files filtered to remove inward ligations (see below). Loops were called at 5 kb resolution using the juicer hiccups tool with KR normalization. CTCF loops were identified with juicer motifs using publicly available SMC3 (ENCSR000EGW), RAD1 (ENCSR000FAD), and CTCF (ENCSR000EGM) ChIP-seq in K562^52^. Differential loops were found by calling loops on biological replicates, finding reproducible loops within conditions, and then comparing the consensus loop lists between conditions. Aggregate peak analysis was performed using the juicer apa tool.

#### Short-range contact probability

Contact probability is calculated from pairs files by first separating reads based on ligation orientation, subtracting positions of cis pairs to find contact distance, then plotting a histogram of those contact distances. It is necessary to separate pairs by ligation orientation because contact probability curves for each orientation are shifted relative to each other – this is due to the fact that read positions will correspond to the entry or exit of each nucleosome in a pair depending on their ligation orientation, and accordingly, the contact distance will include or not the lengths of the nucleosomes. Pairs are separated by ligation orientation based on the strand information as follows: +/- pairs were designated “inward” ligations, -/+ pairs were designated “outward” ligations, and +/+ or -/- pairs were designated “tandem” ligations. To look at short-range contact probability in specific genomic regions, bam files were first intersected with bed regions using bedtools intersect (v 2.30.0) (cite), then pairs with read IDs matching reads in the intersected bam were used for contact distance calculation.

Nucleosome contact peaks are called from the contact probability curves by finding where the second derivative of the curve is negative. NRL is calculated from these peaks by finding the average basepair distance between maxima and N/N+odd:N/N+even ratios of nucleosome contacts are calculated by summing N/N+3 and N/N+5 contact probabilities, summing N/N+2 and N/N+4 contact probabilities, and diving the two values. A higher ratio indicates an enrichment of N/N+odd contacts.

### RNA-seq analysis

RNA-seq data was analyzed by first extracting UMIs and filtering for unique reads using fastp (v 0.24.0) (cite) with parameters --umi_loc per_read --umi_skip 2 --umi_len 5 to match the UMI scheme used by Genewiz. The UMI-filtered data was then aligned to an index composed of combined GRCh38 and ERCC transcriptomes using STAR alignment (v 2.7.11b) (cite) (parameters: --outFilterType BySJout --outFilterMultimapNmax 15 --alignSJoverhangMin 8 --alignSJDBoverhangMin 1 --outFilterMismatchNmax 500 –outFilterMismatchNoverReadLmax 0.05 --alignIntronMin 20 --alignIntronMax 1000000). A counts matrix of paired-end fragments over genes was made from the aligned reads using featureCounts (subread v 2.0.6) (Liao Y, Smyth GK and Shi W (2014)) and the combined gencode.v38-ERCC transcriptome, then DESeq2 (v 1.42.0) was run on genes with at least 10 total counts, using ERCC genes as the control gene set with estimateSizeFactors.

### ATAC-seq data processing

ATAC-seq data was aligned using Bowtie2^53^ filtered, and shift-corrected using deeptools alignmentSieve parameter, which shifts the plus strand by +4 and the minus strand by –5 to account for the Tn5 homodimer which leaves 9bp of DNA between the two Tn5 molecules. Peaks were called using macs2^54,55^ reproducible peaks were found between replicates using Irreproducible discover rate (IDR)^56^ , and a master peak set was made by listing reproducible peaks from both conditions and concatenating adjacent peaks. GenomicRanges,, an R package, is used to load the master peak for further downstream analysis in R. Differential peak analysis was carried out with DESeq2 separately on peaks within 5 kb of a TSS and those further than 5 kb from any TSS. Fragment length distributions were found from filtered bam files using samtools^57^.

Fragment length distributions (FLD) were plotted over ATAC-seq FLD bed files that intersected K562 ChromHmm regions. To get bed files that consisted of accurate fragment lengths, we made ATAC-seq bams that had replicates merged by condition using samtools, then bedpe files were created using bedtools bamtobed with the parameter bedpe. Finally, using awk to get the chromosome name, forward read start coordinates and reverse read end coordinates, which represents the true length of a fragment read and saved that to a bed file. Next, to get the fragments that intersected chromHmm regions, we used bedtools intersect to find and record only the ATAC-seq true fragments that overlapped any of the chromHmm regions. This represents our ATAC-seq fragments found in chromHmm regions bed file.

### CUT&Tag data processing

CUT&Tag data was aligned with BWA mem^58^, filtered, with duplicates removed. Peaks were called using Sicer2^59^, then concatenated the peak set from each replicate from both conditions to create a master peak set.

Bigwig files were created using deeptools^60^ bamCoverage with reads extended and CPM normalized.

Differential peak analysis was done with DESeq2^61^ using the master peak set as regions for aligned reads to be tallied over. Up and down peaks are those that pass two thresholds; 1) Padj value < 0.05, and 2) Log2fold change > |2|. Principle component analysis (PCA) plots were generated using DESeq2 results to show the variance between conditions and replicates to ensure unwanted batch effects were not playing a pivotal role underlying the data.

ChIPseeker^62,63^ is used to annotate the differential peaks. This gives insights into the distribution of peaks in various regions of the genome, as seen in the legend of the plot labeled features.

Heat maps and profile plots were generated using deeptools. To show the reliability of peaks called, we took the differential peaks and plotted the bigwig signal over the center of said differential peaks with 5kb up and down stream. This region illustrates the difference in signal between the two conditions H1low and Scrambled. We also plotted the Cut&Tag H3K27me3 signal over Pro-seq nascent transcription regions to discover if H1 linker histone affecting compaction plays a pivotal role in nascent transcription. Another profile/heatmap plot generated by our pipeline shows our Cut&Tag H3K27me3 signal over differential ATAC-seq peaks. The interaction between chromatin compaction state and loss of H1 illustrates that as we lose linker histone, there is more accessibility.

### Nextflow pipeline data processing

Nextflow^64^ is used to create a reproducible and scalable pipeline that incorporates many of the tools mentioned in the methods section. We have two pipelines engineered to handle epigenomic sequencing techniques. The first Risca Lab pipeline (NEXDEP) can process fastq reads from ATAC-seq, Cut&Tag, Cut&Run, ChIP-seq assays to align, filter, give quality control metrics and preprocess data to produce bam files with sequence alignment information. The second Risca Lab pipeline was engineered specifically to call peaks from previously mentioned assays and provide downstream analysis and plots such as heatmaps, MA-plots, PCA plots, and peak annotation information, along with many other custom analytical Nextflow workflow techniques that partially aided in completion of this study.

### IDA and data analyses

IDA was performed to generate reference spectral libraries for SWATH data quantification. The IDA method was set up with a 200 ms TOF-MS scan from 300 to 1,250 Da, followed by MS/MS scans in a high-sensitivity mode from 100 to 1,500 Da of the top 25 precursor ions above 100 cps threshold (80 ms accumulation time, 100 ppm mass tolerance, rolling collision energy, and dynamic accumula-tion) for charge states (z) from +2 to +5. IDA files were searched using ProteinPilot (version 5.0.2, ABSciex) with a default setting for tryptic digest and IAA alkylation against a protein sequence data-base.

The Homo sapiens proteome FASTA file (82,493 protein entries, UniProt UP000005640) augmented with sequences for common contaminants was used as a reference for the search. Up to two missed cleavage sites were allowed. Mass tolerance for precursor and fragment ions was set to 100 ppm. A false discovery rate (FDR) of 5% was used as the cutoff for peptide identification.

### SWATH acquisitions and data analyses

For SWATH (SWATH-MS, Sequential Window Acquisition of All Theoretical Mass Spectra) acquisitions (Zhu et al., 2014), one 50-ms TOF-MS scan from 300 to 1,250 Da was performed, followed by MS/MS scans in a high-sensitivity mode from 100 to 1,500 Da (15 ms accumulation time, 100 ppm mass tolerance, +2 to +5 z, rolling collision energy) with a variable-width SWATH window (Zhang et al., 2015). DIA data were quantified using PeakView (version 2.2.0.11391, ABSciex) with SWATH Ac-quisition MicroApp (version 2.0.1.2133, ABSciex) against selected spectral libraries generated in Pro-tein-Pilot. Retention times for individual SWATH acquisitions were calibrated using 25 or more pep-tides for plectin (PLEC, UniProt Q15149) and myosin-9 (MYH9, UniProt P35579), two abundant pro-teins that were highly representative in the IDA ion library and all SWATH acquisitions. The following software settings were utilized: up to 25 peptides per protein, 6 transitions per peptide, 95% peptide confidence threshold, 5% FDR for peptides, XIC extraction window 10 minutes, and XIC width 100 ppm.

